# A survey of vaccine-induced measles IgG antibody titer and the verification of changes in temporal differences of measles vaccination in young adults

**DOI:** 10.1101/622613

**Authors:** Hiraku Sasaki, Tomoko Fukunaga, Ai Asano, Yoshio Suzuki, Yuko Nakanishi, Junzi Kondo, Hiroki Ishikawa, Nobuto Shibata

## Abstract

In Japan, sporadic measles cases increased rapidly in 2019 compared to that in past 6 years. To clarify the persistence of immunity against measles in young adult, this study explored the prevalence of IgG antibody titer against measles virus in 18-to 24-year-old young adult participants. Measles-specific IgG antibody titers determined by enzyme immunoassay in serum samples collected from 506 participants between 18 to 24 years were assessed through statistical analyses. Multiple regression analysis revealed that the distribution of measles IgG antibody titers was significantly correlated with medical history (*P* < 0.05), while there was no significant difference among the number of vaccinations related to measles IgG titers. Further, measles IgG titers were significantly different, which was determined by the temporal change that elapsed period after last vaccination (*P* < 0.05). These results indicate that periodic vaccination against measles is required for young and older adults to prevent even sporadic measles infection.

## Introduction

In Japan, more than 300 cases of measles were reported, which has increased rapidly compared to that in the past 6 years [1]. Measles cases were reported in 21 of 47 prefectures, and in particular, a third part of measles case were found in Osaka prefecture within 1^st^ to 10^th^ week in 2019 [1]. Taking measles vaccination twice in lifetime induced acquired immunity against measles in more than 95% of people [2, 3]. For the measles vaccination program in Japan, a person born after April 2006 is defined to have received 2 doses of measles vaccine as a periodic inoculation at 1-2 year and 5-7 years of age [4]. Before the Application of these regulations, there used to be only primary vaccination after 12 months of age. Thereafter, there was a change in the regulations of the Ministry of Health, Labor and Welfare recommending 2 doses of measles vaccination for people born before April 2006 with secondary inoculation being received within the duration of 5 years (2008–2012, equivalent to 13 and 18 years of age) as special measures for periodic inoculation [4]. Within the 5-years, the rate of vaccination ranged from 77.3 to 94.3% [5]; however, recent vaccination rate exhibited more than 90% of people receiving measles vaccines twice in Japan [6]. Except for the people who received secondary vaccination at limited periods, almost all young adults born after April 2006 were vaccinated twice before they were 7 years of age. A large measles outbreak is thought to be physically impossible in later childhood due to the implementation of the measles vaccine program; however, there was insufficient knowledge about the existence of protective antibody titer against measles in young adults who received secondary measles vaccine within the 5 years duration limit.

Currently, measles cases are based on age bracket. In brief, more than 50% of measles cases were seen in 15–29-year-old individuals, while less than 10–14-year-old constituted only 12% [1]. Almost all young adults around 20 years of age were subjected to the special measures of periodic inoculation but all of them did not receive the same manner of measles vaccination. In particular, temporal changes in protective serum IgG antibody against measles virus may vary in spite of receiving secondary vaccination lately compared to young adults. However, temporal changes in measles IgG antibody titer in generation of young adults that was subjected to special measures of measles vaccine are obscure. Thus, there is a need to elucidate the factors involved in the risk of the infection as well as the ways of reducing the occurrence of the infection.

To clarify the persistence of immunity against measles in young adults, this study explored the prevalence of IgG antibody titer against measles virus in 18–24-year-old young adult participants, and determined the relationships between temporal changes in the antibody.

## Materials and Methods

### Study design

The surveillance was carried out by obtaining samples of sera from 506 young adults between 18 to 24 years of age, who were first-year students in Juntendo University to assess the prevalence of specific IgG antibodies against measles virus. The serum samples were obtained from January 2018 to April 2018. Simultaneously, vaccine history was collected from each individual’s maternity passbook. The studied participants also filled out questionnaires aimed at obtaining information about the medical history of vaccination against measles and natural measles infection. The collected information was used for proper interpretation of the obtained results.

The study protocol was approved by the Ethics Committee of Faculty of Health and Sports Science, Juntendo University (number 30-2) and informed consent was obtained from all participants and their parents before the study.

### Laboratory methods

To determine measles-specific serum IgG antibody titer, the enzyme immunoassay (EIA) was performed in the laboratory of BML, INC. (Tokyo, Japan) using a measles virus immunoglobulin test kit (the Measles IgG-EIA manufactured by Denka Seiken., Co., Ltd, Tokyo, Japan). For the virus antigen, Toyoshima strain was used.

### Statistical analysis

To identify the factors that influence measles-specific IgG antibody titer, multiple regression analysis was performed. To avoid multicollinearity, factors were preliminary determined with bivariate analysis and were confirmed to have no correlation. Differences in the IgG antibody titers between the participants that had a medical history of measles and those without history were evaluated with the unpaired *t*-tests. To compare differences in IgG antibody titers among the numbers of vaccination and temporal characteristics, a one-way analysis of variance (one-way ANOVA) was employed. When there were significant statistical differences, the data were further analyzed using the Bonferroni *post hoc* test to determine the significance between the groups. Differences were considered significant for *P* values of less than 0.05.

## Results

### Outline of survey participants

During the blood collection for determining antibody titers in this surveillance, participants who were 18 years of age were 80% of total survey participants while the remaining 20% of participants were ≥ 19 years old. All the subjects who were 19 years old did not receive secondary inoculations. Approximately 80% of the participants received secondary vaccination, and 10% received only primary vaccination (Table 1). Approximately 4% of participants had a medical history of measles based on self-assessment information.

**Table 1.**
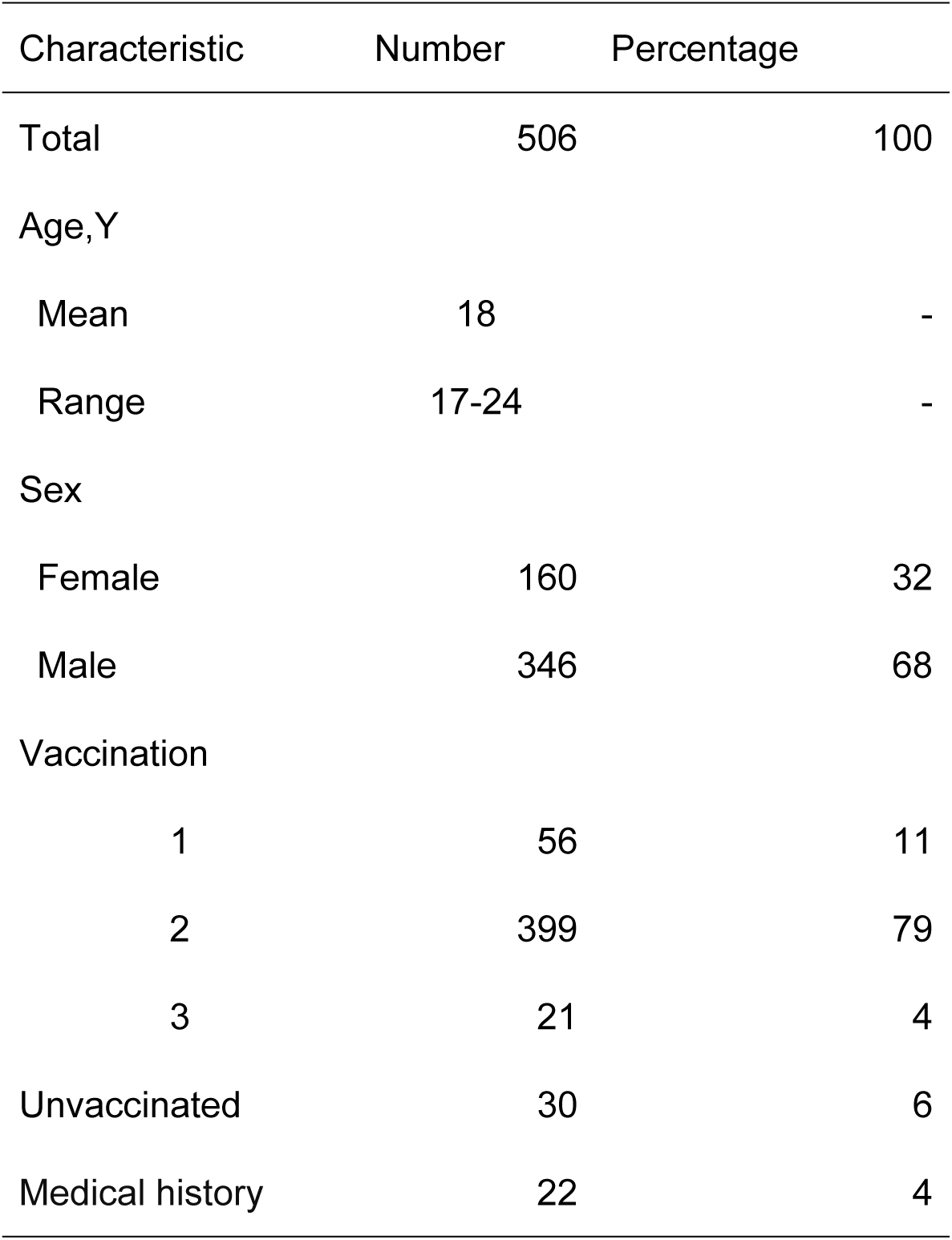
Outline of measles-vaccine induced antibody survey participants

| Characteristic | Number | Percentage |
| --- | --- | --- |
| Total | 506 | 100 |
| Age, Y |  |  |
| Mean | 18 | - |
| Range | 17-24 | - |
| Sex |  |  |
| Female | 160 | 32 |
| Male | 346 | 68 |
| Vaccination |  |  |
| 1 | 56 | 11 |
| 2 | 399 | 79 |
| 3 | 21 | 4 |
| Unvaccinated | 30 | 6 |
| Medical history | 22 | 4 |

### Measles-IgG titer and temporal characterization

The mean value of measles-specific IgG antibody titer determined by EIA exhibited 13.4 ±12.0 (Mean ± standard deviation) ranging from 0.8 to 128 (Table 2). The period of the month when primary and secondary vaccines were received exhibited 23.6±32.5 and 152.2 ±25.4 month, respectively. The recommended age for primary measles vaccination was to be from 12 to 90 months during 1995 to 2000, and therefore, the duration for primary vaccination was extended. Almost all participants corresponded to the special measure that was recommended for secondary vaccination at 13 years of age during 2008 to 2013, indicating that more than 80% of participants received secondary vaccination at the assigned period. Regarding all the vaccinated participants, an average of 7 years passed from last vaccination.

**Table 2.**
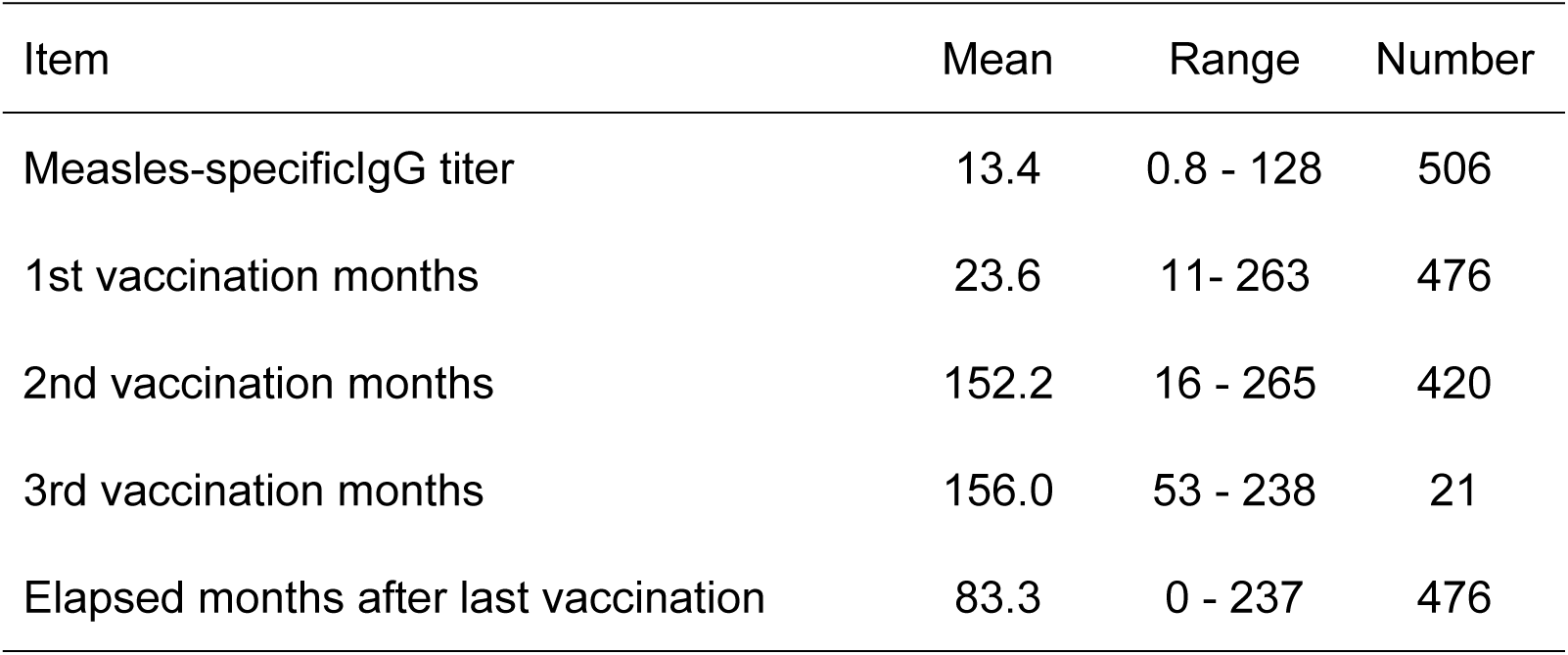
Measles-specific IgG antibody titer and temporal characteristics of participants.

### Divergence of measles-IgG antibody titers by medical history

To discern mode of measles IgG antibody titer’s distribution, multiple regression analysis was performed for the IgG antibody titers and temporal characteristics. It was observed that almost all parameters related to temporal characteristics had multicollinearity. Thus, multiple regression analysis was performed based on Table 1 listed items devoid of temporal information. Of these, the distribution of measles IgG antibody titers was significantly correlated with medical history of measles (*P* < 0.05). Then, the differences in measles IgG antibody titers between presence and absence of medical history were compared by the unpaired *t*-tests (Figure 1). The IgG antibody titers collected from participants who had medical history of measles (27.0 ± 31.8) were significantly higher than titers from participants who had no history (12.8 ± 9.9, *P* < 0.05).

**Figure 1.**
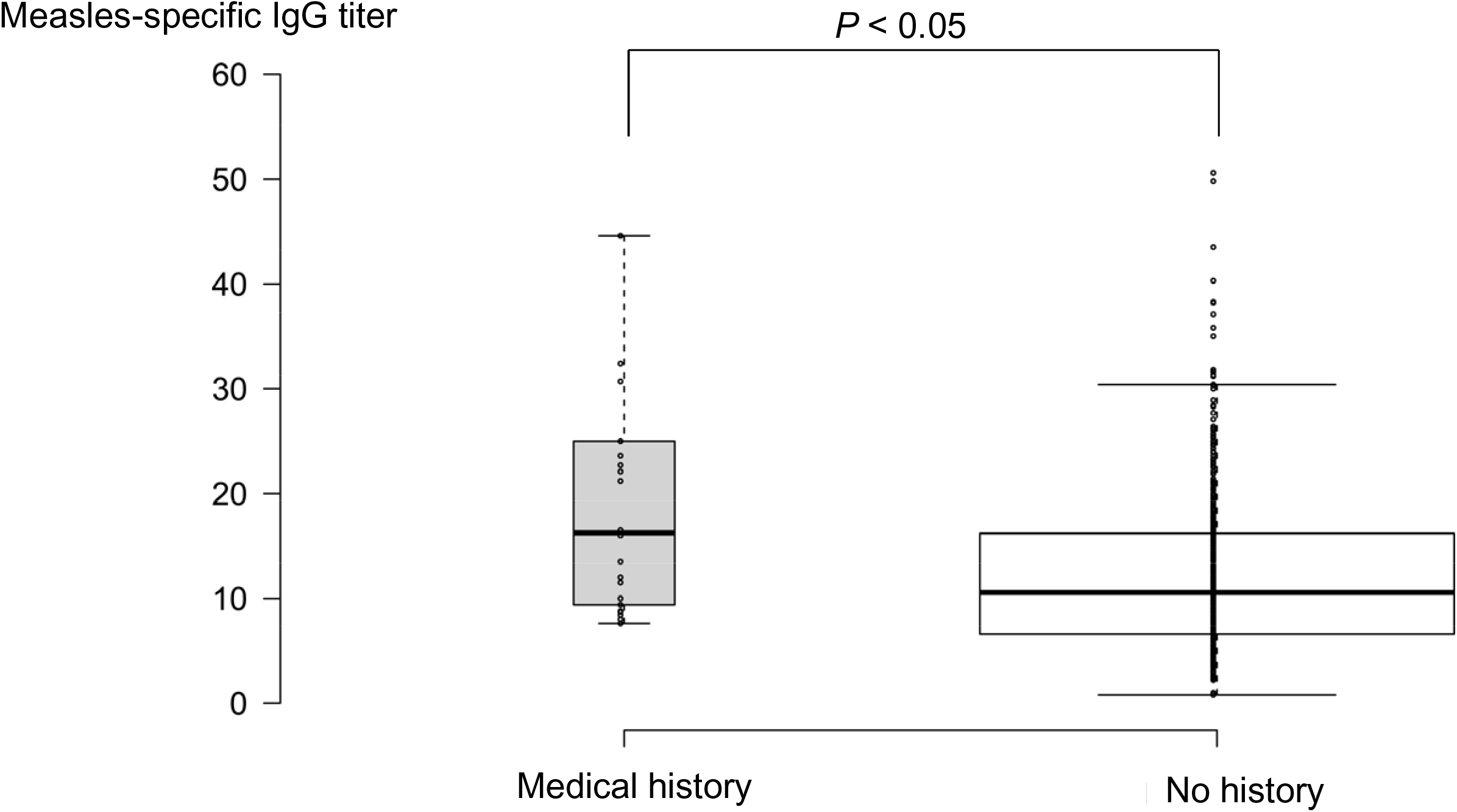
Comparison of measles-specific IgG antibody titers based on the medical history of participants. The plots of IgG antibody titer that were more than 60 were omitted from Figure 1. The whiskers extended to data points that were less than 1.5 x IQR away from 1st/3rd quartile.

Figure 2 shows measles IgG antibody titers that were divided into each numbers of measles vaccinated and unvaccinated participants. The measles IgG antibody titers from the participants who had a medical history of measles were excluded from Figure 2 and statistical analysis since almost all participants who had a medical history had been unvaccinated and exhibited higher IgG antibody titer than participants who had no history. The results of one-way ANOVA showed that there was no significant difference among the number of vaccinations.

**Figure 2.**
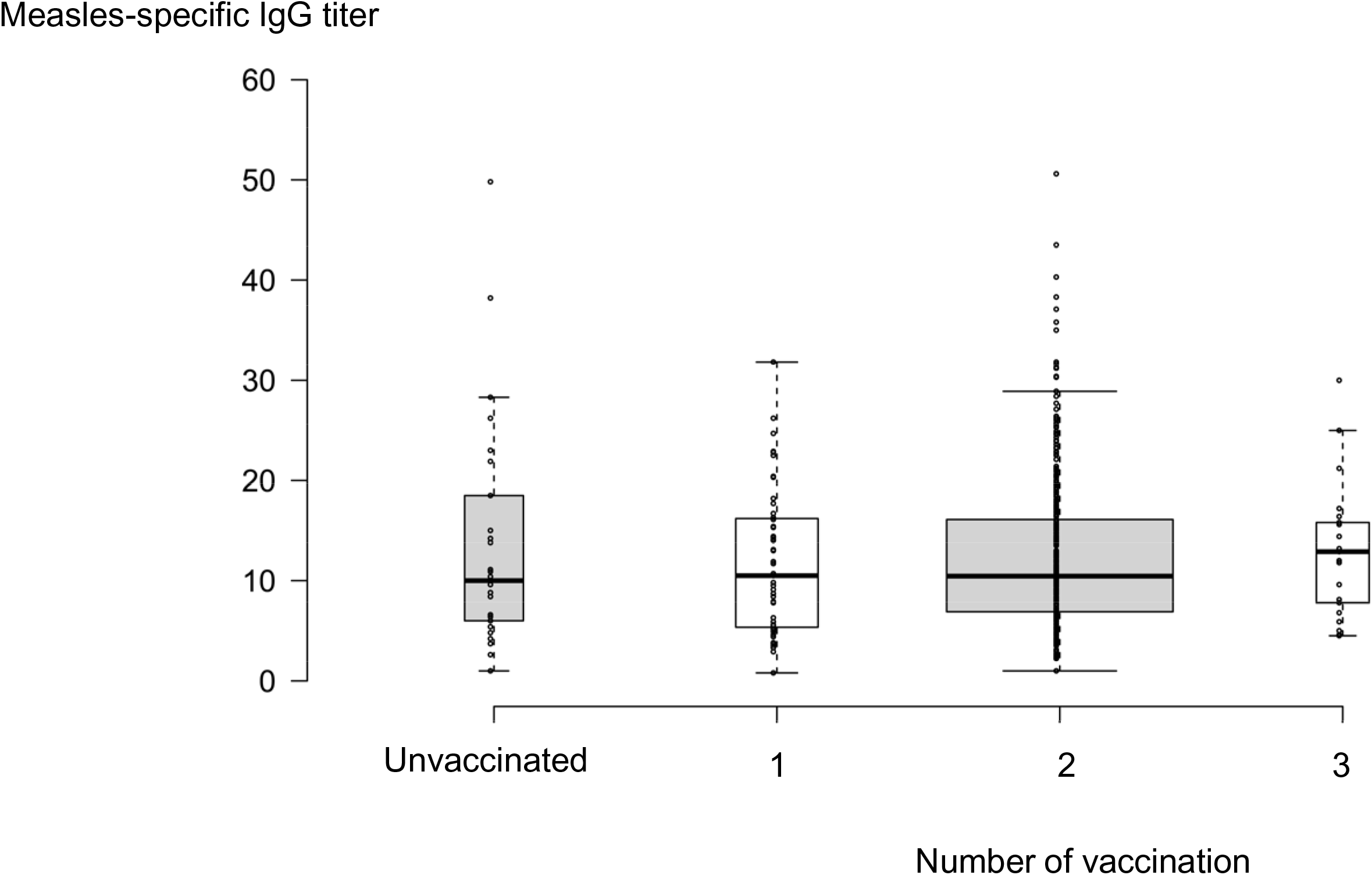
Comparison of measles-specific IgG antibody titers between the number of vaccinated and unvaccinated participants. The plots of IgG titer that were more than 60 were omitted from Figure 2. The measles IgG antibody titers from the participants who had a medical history were excluded from Figure 2 and statistical analysis. The whiskers extended to data points that were less than 1.5 x IQR away from 1st/3rd quartile. There were no significant differences between number of vaccinated and unvaccinated participants based on one-way ANOVA (P ≥ 0.05).

### Temporal changes in measles-IgG antibody titers

To verify the temporal changes in measles IgG antibody titers after the last vaccination, the elapsed periods were divided into 5 periods including 3 month or less (n = 9), 4–12 months (n = 8), 13–60 months (n = 31), 61–72 months (n = 279), and 73 months or more (n = 73). In measles vaccine program, almost all participants received secondary vaccination at 13 years of age, and thus, approximately 70% of participants were observed in the period 61–72 months. Excluding measles IgG antibody titers from the participants who had a medical history and those who were unvaccinated, one-way ANOVA showed that there were significant differences among the 5-periods (Figure 3, *P* < 0.05). Although secondary period (4-12 month) contained a few dispersed measles IgG antibody titer values, the mean values of measles IgG antibody titer decreased with the prolonged period. The Bonferroni *post hoc* test showed that there were 4 significant differences between each period (*P* < 0.05), indicating that measles IgG antibody titer retains high value if elapsed period is short after last vaccination.

**Figure 3.**
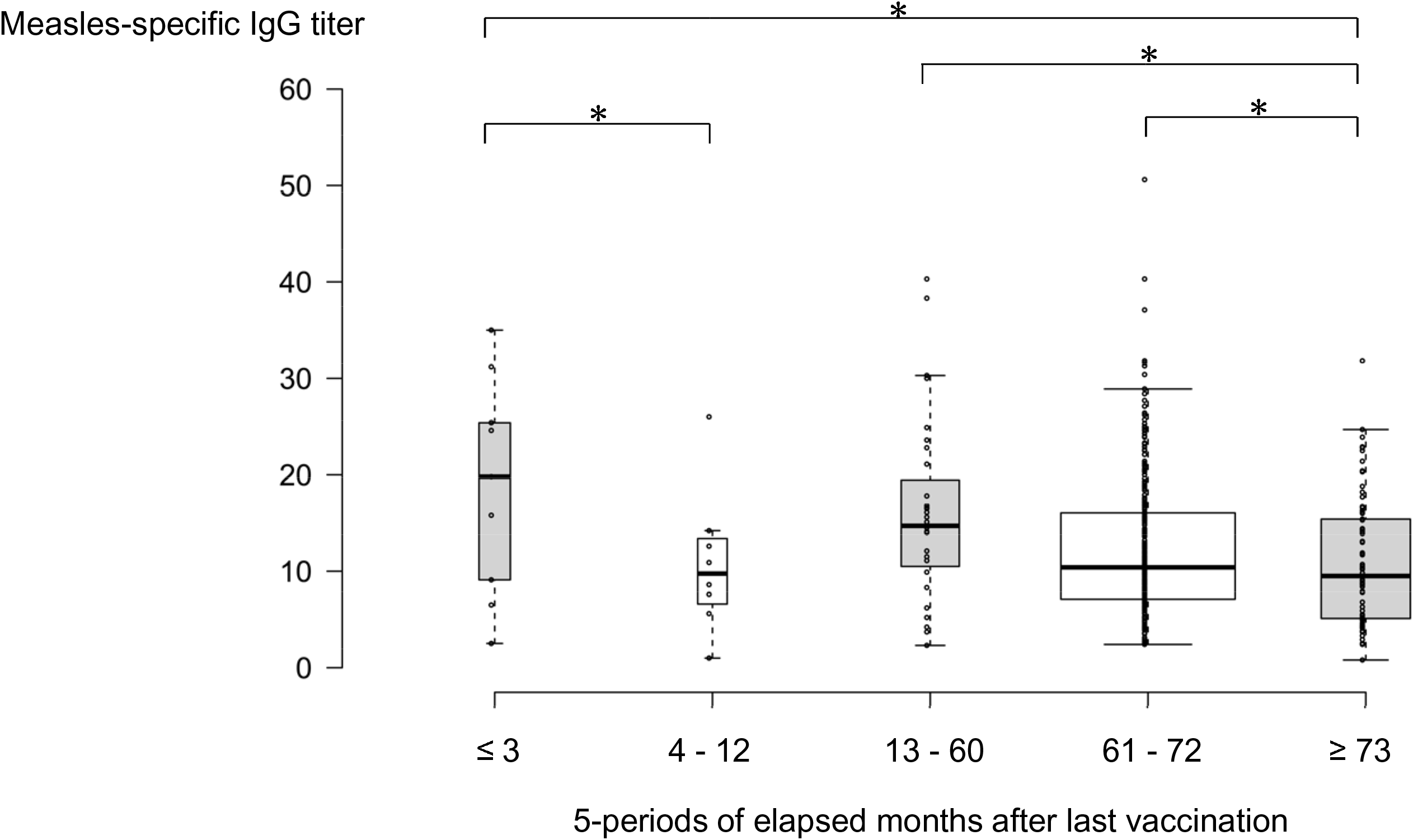
Comparison of measles-specific IgG antibody titer among 5-periods of elapsed months after last vaccination The plots of IgG titer that were more than 60 were omitted from Figure 3. The measles IgG antibody titers of both the participants who had a medical history and the unvaccinated participants were excluded from Figure 3 and statistical analysis. The whiskers extended to data points that were less than 1.5 x IQR away from 1st/3rd quartile. One-way ANOVA showed that there were significant differences among the 5-periods (P < 0.05). An asterisk denotes a significant difference between periods with connected line (P < 0.05).

## Discussion

To protect from measles virus infection, measles-specific serum IgG antibody titer determined by the EIA method was required 12 or more, and in case of unsatisfied titer, additional vaccination was recommended [7]. Although average measles IgG antibody titer of total participants slightly exceeded 12, 56% of measles IgG antibody titers from 284 participants showed less than 12. These results indicate that more than half of the studied participants required additional dose to protect against measles. According to WHO announcement, Japan was verified as having achieved measles elimination that was defined as interruption of endemic measles virus transmission for at least 36 months [8]. Nevertheless, measles cases from foreigners in Japan have occurred sporadically [1, 9-12]. Based on our surveillance, one of the causes may be non-persistent protection by measles IgG antibody titer in young adults. When measles was endemic in Japan, approximately 200,000 people, mainly children, were infected with measles virus in the year 2000 [13, 14]. Thereafter, an attempt to raise coverage of measles vaccination and to enforce 2 doses vaccination rigorously was successfully achieved, which led to the elimination-period [8]. Currently, protection against measles infection has been achieved; “however, the generation with insufficient immunization against measles virus is mainly the young adults suggesting that periodic monitoring of measles epidemic and acquired immunity against measles virus in young adult is required.

Further, this surveillance focused on the temporal status of measles IgG antibody from last vaccination, although there was no significant difference in measles IgG antibody titers after primary and secondary vaccination. In the United States, vaccination schedule and patterns are similar to those in Japan, and measles was declared eliminated with the absence of continuous measles transmission for a period greater than 12 months in the United States in the year 2000 [15]. However, endemic outbreaks of measles were reported to be yet to occur [16-18], and in rarity, a part of those outbreaks was caused by unvaccinated population [19]. For primary vaccine failure, CD46 and TLR8 variants were considered involved in the occurrence of measles vaccine failure [20]. Although it will be possible to have such cases, epidemiological studies have demonstrated the efficacy of measles vaccine. In brief, 95% of children who received measles vaccine acquired immunity against measles virus, and further, additional secondary vaccine led to more than 99% immunization in children [21-23]. These results revealed that epidemic outbreaks might be caused by the unvaccinated population or large number of international travelers [3]. Further, according to the large surveillance by healthcare workers, serum measles IgG antibody titers from adults less than 29 years of age showed susceptibility to measles [24]. Single dose of measles vaccine in adults was reported to have significantly increased serum IgG titers even in the initial insufficient IgG titers [25]. Sporadic infection may occur in young and older adults under unprotected conditions and with reduced vaccine-induced IgG antibody titer due to the temporal changes after last vaccination and consequent susceptibility to infection. These results also suggest that preventive vaccination against measles is required for young and older adults to prevent even sporadic measles cases.

## Acknowledgements

This work was partially supported by JSPS KAKENHI Grant Number 16K07095.

